# The Future of *Juniperus procera* in Ethiopia under Climate Change Scenarios - Implications for Ecosystem Rehabilitation and Biodiversity Conservation

**DOI:** 10.1101/2025.06.25.661646

**Authors:** Zerihun Woldu, Gete Zeleke

## Abstract

*Juniperus procera*, a keystone conifer of Ethiopia’s dry Afromontane forests, is increasingly threatened by climate change and land-use pressures. This study employed an ensemble species distribution modeling (SDM) approach integrating Random Forest, MaxEnt, and Boosted Regression Trees to predict the species’ current and future habitat suitability under two climate scenarios: SSP2-4.5 (intermediate emissions) and SSP5-8.5 (high emissions) for the 2061-2080 period. High-resolution bioclimatic variables and a curated dataset of georeferenced occurrence records were used to calibrate and validate the models. To ensure robustness, variable selection was performed through correlation filtering, and models were evaluated using AUC, TSS, and Kappa metrics.

Model outputs revealed a marked contraction of highly suitable habitats under both scenarios, with losses reaching up to 85% under SSP5–8.5. In contrast, areas currently classified as unsuitable are projected to gain suitability, particularly in lower elevation belts of moist Afromontane zones, and higher elevation belts in northern Ethiopia suggesting potential downslope range shifts. Delta maps and transition matrices indicated widespread redistribution, including fragmentation and quality downgrading of core habitats.

These findings highlight the dual challenges and opportunities posed by climate change for *J. procera*. While habitat loss and ecological fragmentation threaten long-term species viability, gains in new areas may offer pathways for adaptation provided dispersal, land use, and landscape connectivity are addressed.

This study underscores the need for climate-informed, spatially explicit conservation strategies. Recommendations include prioritizing in-situ protection of climate refugia, promoting assisted migration to emerging suitable zones, and restoring degraded landscapes to support ecological resilience. These insights support ecosystem-based adaptation and sustainable forest management planning under Ethiopia’s national conservation and afforestation initiatives.

## 1. Introduction

The dry evergreen Afromontane forests of Ethiopia are among the most ecologically significant terrestrial ecosystems in the Horn of Africa, harboring a high degree of endemism and biological diversity. These forests are also of immense socioeconomic importance, providing essential ecosystem services such as water regulation, carbon storage, erosion control, and a range of forest products that support rural livelihoods. However, over the past several decades, the extent and integrity of Ethiopia’s Afromontane forests have been severely compromised by a suite of anthropogenic pressures. Rapid deforestation, largely driven by agricultural expansion, fuelwood collection, settlement growth, and unsustainable land management practices, has led to the fragmentation and degradation of forest habitats (Gete Zeleke & Hurni, 2001; Hurni et al., 2010). Superimposed on these direct human drivers is the growing threat of climate variability and long-term climate change, which is altering temperature and precipitation patterns across Ethiopia’s highlands.

Among the dominant species in these forests, *J*.*procera* (African pencil cedar) plays a keystone role in maintaining ecosystem structure and function across the mid-to high-elevation zones of the Ethiopian highlands (Negash, 2010). The keystone species concept, introduced by Paine (1969), originally referred to marine predators but now includes structural species like *J. procera*, whose ecological roles far exceed their abundance in maintaining system stability and resilience. As a long-lived conifer with broad ecological amplitude and cultural significance, *J. procera* contributes to soil stabilization, microclimate regulation, biodiversity support, and the maintenance of hydrological balance, particularly in montane catchments feeding the upper Nile Basin (Zerihun Woldu, 1999).

Despite its importance, *J. procera*’s distribution has contracted significantly due to anthropogenic pressures and climatic shifts (Aerts et al., 2006; Tolessa, Senbeta & Kidane, 2017). This decline threatens forest biodiversity and critical ecosystem services such as watershed stability and baseflow in downstream rivers including the Nile (Bewket & Sterk, 2005). Understanding the current and future potential distribution of *J. procera* is therefore essential for guiding conservation, reforestation, and climate adaptation efforts.

In response to these multifaceted threats, predictive tools such as Species Distribution Models (SDMs) have become increasingly indispensable for assessing the potential impacts of environmental change on species’ geographic ranges. SDMs combine known species occurrence records with environmental variables such as climate, topography, and land cover—to estimate habitat suitability across space and time. These models offer valuable insights for anticipating range contractions, expansions, or shifts under various climate change scenarios, thereby supporting proactive conservation and management strategies. For long-lived and ecologically pivotal species like *J*.*procera*, which may be particularly sensitive to changing climatic conditions, SDMs can provide critical evidence for identifying climate refugia, restoration priority zones, and areas vulnerable to habitat loss.

In this study, we employ an ensemble SDM approach that integrates multiple algorithms including Random Forest (RF), Maximum Entropy (MaxEnt), and Boosted Regression Trees (BRT) to improve prediction robustness and reduce model uncertainty. By projecting the current and future distribution of *J. procera* under two Shared Socioeconomic Pathways (SSP2-4.5 and SSP5-8.5), we aim to elucidate potential changes in habitat suitability across Ethiopia’s highlands and inform adaptive forest management in the face of a changing climate. This modeling aligns with Ethiopia’s Green Legacy Initiative, a national campaign to restore degraded lands, combat desertification, and enhance carbon storage through large-scale tree planting (FDRE, 2019). Integrating ensemble SDM results into restoration planning can target climatically suitable areas where *J. procera* populations are likely to persist or expand. Prioritizing afforestation in these zones can increase survival rates, maximize ecological benefits, and support recovery of degraded ecosystems. Consequently, spatial predictions of suitable habitats provide critical guidance to align restoration with climate resilience and biodiversity conservation goals.

While local conservation efforts are vital, the challenges facing *J. procera* reflect broader global patterns of biodiversity loss driven by climate change and habitat alteration. Changes in temperature and rainfall disrupt habitats and ecological relationships, pushing many species toward extinction (Thomas et al., 2004). These pressures are often exacerbated by land use and land cover changes such as deforestation and agricultural expansion, which fragment habitats and hinder species’ ability to adapt or migrate (Foley et al., 2005; Newbold et al., 2015).

The overarching goal of this study is to assess the current and future habitat suitability of *J*.*procera* across Ethiopia under changing climatic conditions. Specifically, we aim to: (i) model the current distribution of suitable habitats for *J. procera* using a robust ensemble of SDM algorithms; (ii) project potential shifts in habitat suitability under mid-century climate scenarios (SSP2-4.5 and SSP5-8.5); (iii) quantify the extent and spatial pattern of gains, losses, and stable areas in *J. procera* habitat; (iv) identify priority areas for conservation, restoration, or assisted migration; and (v) provide spatially explicit guidance to support Ethiopia’s Green Legacy Initiative and inform restoration, watershed management, and broader conservation strategies.

By integrating high-resolution bioclimatic data with advanced modeling techniques, this study provides a comprehensive evaluation of the vulnerability and resilience of *J. procera* in the face of climate change. The findings are intended to inform landscape-scale conservation planning and to support national reforestation and ecosystem restoration initiatives, such as Ethiopia’s Green Legacy program.

## 2. Literature Review

One of the earliest comprehensive vegetation classifications in East Africa was by Pichi-Sermolli (1957), who mapped major vegetation types including Ethiopia’s dry evergreen montane forests dominated by *J*.*procera*. This seminal work established the ecological significance of *J. procera* in the Ethiopian highlands, influencing subsequent ecological and botanical research.

Friis et al. (2010) expanded on this foundation through a detailed vegetation atlas of Ethiopia, delineating 15 major vegetation types. The “Dry evergreen Afromontane forest and grassland complex” was highlighted, with *J. procera* identified as a dominant canopy species in the dry single-dominant Afromontane forest subtype. This atlas, integrating field observations and GIS analyses, refined the understanding of the species’ ecological niches and spatial distribution patterns.

*J. procera* occurs predominantly between 1,500 and 3,200 meters above sea level, thriving where moderate temperature and moisture conditions prevail (Friis et al., 2010). Its lower altitudinal range is limited by aridity, elevated temperatures, and human disturbances such as agriculture and grazing, while low temperatures and frost constrain its upper range (Zerihun Woldu, 1999). The species coexists with Afromontane associates including *Olea europaea* subsp. *cuspidata, Podocarpus falcatus*, and *Hagenia abyssinica* across transitional forest belts (Zerihun Woldu, 1985). Abiotic factors such as precipitation, temperature, and soil moisture are critical determinants of *J. procera*’s habitat suitability, with a preference for well-drained soils typical of mid-elevation refugia (Friis et al., 2010; Teketay, 1992).

As a long-lived canopy conifer, *J. procera* contributes substantially to ecosystem structure and function. It stabilizes soils, regulates microclimates, and supports biodiversity by providing resources for frugivorous fauna, facilitating seed dispersal and maintaining ecological connectivity (Negash, 2010; Borghesio et al., 2004). The decline of *J. procera* risks disrupting trophic interactions and forest integrity at landscape scales.

Socioeconomically, *J. procera* has considerable value. Its durable timber is employed in religious and traditional architecture, its essential oils have commercial applications, and its plant parts are widely used in traditional medicine (Adams, 1990; Abebe et al., 2003). Thus, its conservation carries both ecological and cultural importance.

A significant climate-driven phenomenon in Ethiopian montane systems is the upslope shift of C_4_ grasses into zones formerly dominated by C_3_ species (Zerihun Woldu, 1991). This encroachment, driven by rising temperatures and moisture stress, favors C_4_ plants such as *Andropogon abyssinicus, Hyparrhenia hirta*, and *Chloris gayana* due to their physiological advantages in water-use efficiency and thermal tolerance (Sage et al., 1999; Taylor et al., 2014). Comparable patterns are documented in tropical montane ecosystems globally, highlighting widespread altitudinal vegetation shifts under climate change (Liu et al., 2018; D’Antonio et al., 2000).

Species Distribution Models (SDMs) are vital tools for spatially explicit predictions of species ranges by correlating occurrence data with environmental predictors. Ensemble modeling approaches, combining MaxEnt, Random Forest, and Boosted Regression Trees (BRT), enhance predictive accuracy and reduce uncertainty (Araújo & New, 2007; Thuiller et al., 2009). For long-lived and structurally important species like *J. procera*, ensemble SDMs provide robust projections of habitat suitability under current and future climates, capturing landscape-level variability (Franklin, 2013; Morán-Ordóñez et al., 2017).

In data-limited but biodiversity-rich regions such as Ethiopia, ensemble SDMs improve confidence in delineating climate refugia, range shifts, and contraction areas critical for landscape-scale conservation planning. They support adaptive management under multiple emission scenarios (e.g., SSP2-4.5 and SSP5-8.5), offering spatially explicit guidance to align conservation with restoration initiatives and policy frameworks (Guisan et al., 2013). Such models underpin strategies for landscape connectivity, ecosystem restoration, and biodiversity targets consistent with the Aichi Biodiversity Targets, the Bonn Challenge, and the UN Decade on Ecosystem Restoration (IPBES, 2019).

However, research applying SDMs to *J. procera* remains sparse and geographically limited. Zenebe et al. (2024) modeled habitat suitability but restricted their scope to remnant forest patches, constraining landscape-level extrapolation. Earlier descriptive work by Kerfoot (1963) lacked spatial precision and quantitative modeling, while Gebrehiwot et al. (2024) analyzed climate trends without integrating spatially explicit projections, limiting actionable conservation insights.

Importantly, few studies explicitly connect SDM outputs with Ethiopia’s Green Legacy Initiative, a large-scale reforestation and restoration campaign, or broader national restoration goals. Addressing this gap requires geographically comprehensive, policy-relevant ensemble SDMs incorporating multiple climate scenarios, which can generate actionable spatial guidance for conservation, restoration, and climate adaptation of *J. procera* across the heterogeneous Ethiopian landscape.

## 2. Methods

### 2.1. Study Area

This study focuses on Ethiopia, located in the Horn of Africa, which spans a wide altitudinal gradient and diverse climatic zones. The Afromontane highlands, ranging from approximately 1,500 to over 3,200 meters above sea level host remnant dry evergreen forests dominated by *J. procera*. These highlands are characterized by temperate conditions, with annual rainfall ranging between 700 and 2,000 mm and mean annual temperatures from 10°C to 20°C (NMA, 2007). The region is ecologically critical and socioeconomically significant, yet increasingly vulnerable to land-use conversion and climatic stressors.

### 2.2. Species Occurrence Data

Presence records of *J*.*procera* were compiled from multiple sources, including the Global Biodiversity Information Facility (GBIF), herbarium databases, and published literature. All records were filtered for spatial accuracy, removing duplicates and those with coordinates falling outside the known native range or in highly disturbed landscapes (e.g., urban centers). The final dataset consisted of 88 georeferenced presence points.

### 2.3. Environmental Variables

To model the species’ ecological niche, we selected a suite of bioclimatic variables from the WorldClim v2.1 database at a 2.5 arc-minute resolution. These included variables relevant to temperature and precipitation seasonality, extremes, and variability. Highly correlated variables (Pearson’s |r| ≥ 0.8) were removed through pairwise correlation analysis, retaining those with greater ecological relevance to coniferous montane species. Additionally, a digital elevation model (DEM) was incorporated to capture the influence of topographic variation.

Future climate projections for the mid-21st century (2061–2080) were obtained for two emission scenarios - SSP2-4.5 (intermediate) and SSP5-8.5 (high) based on CMIP6 models downscaled by WorldClim. All current and future variables were cropped to Ethiopia’s extent and resampled to a consistent resolution using the `terra` package in R. The choice of the 2060–2080 timeframe aligns with the CMIP6 “2070s” projection window, commonly used in species distribution studies due to standardized scenario availability and high-resolution data coverage (IPCC AR6 WG1 Report (2021)). Given *J. procera*’s slow growth and longevity, this temporal horizon is appropriate for capturing ecologically meaningful habitat shifts. Earlier time slices may underestimate cumulative warming and precipitation changes, especially in montane and dryland ecosystems where stressors accumulate gradually (Luedeling et al., 2011). Furthermore, the divergence between moderate and high-emission scenarios becomes more pronounced in this period, enhancing the robustness of scenario-based comparisons.

### 2.4. Species Distribution Modeling

We implemented an ensemble species distribution modeling framework using three widely applied algorithms: Random Forest (RF), Maximum Entropy (MaxEnt), and Boosted Regression Trees (BRT). Model calibration and evaluation were conducted using the `sdm`, `dismo`, `gbm`, and `randomForest` packages in R. Each model was trained using 80% of the presence data and evaluated on the remaining 20%, repeated over 10 replicates for robustness. Pseudo-absence/background points (n = 10,000) were generated randomly within a suitable environmental envelope.

Model performance was assessed using Area Under the Receiver Operating Characteristic Curve (AUC) and True Skill Statistic (TSS). Ensemble predictions were generated using a weighted mean approach based on individual model AUC scores. Suitability maps were reclassified into four categories: None, Low, Medium, and High based on thresholds derived from model sensitivity and specificity trade-offs.

### 2.5. Change Detection and Area Analysis

To assess potential range dynamics, we compared current and future ensemble suitability maps for each scenario. Raster differencing and transition matrices were used to classify areas as lost, gained, stable, or unchanged. All rasters were reprojected into a UTM zone appropriate for Ethiopia (e.g., UTM Zone 37N) to ensure accurate area calculations. Total area and percentage change were computed for each suitability class and change type using the `terra` and `sf` packages. The results were visualized with `ggplot2`, and spatial patterns were evaluated in the context of Ethiopia’s topographic and ecological gradients.

### 2.6. Statistical Analysis

Descriptive statistics were used to quantify the spatial extent and temporal changes in habitat suitability classes under current and future climate scenarios. Specifically, changes in habitat area categorized as gains, losses, or stable zones were computed by overlaying classified suitability maps from the present and future periods. Class-wise transition matrices were generated to capture fine-scale dynamics, enabling the identification of specific suitability shifts such as upgrades (e.g., Low to Medium) or downgrades (e.g., High to None), and to summarize patterns of range expansion, contraction, and persistence.

To test the statistical significance of differences in habitat suitability across climate scenarios and suitability classes, a Generalized Linear Model (GLM) with a Gaussian error distribution was fitted. The GLM evaluated the effects of climate scenario (current, SSP2-4.5, SSP5-8.5) and suitability class (None, Low, Medium, High) on habitat area (in km^2^). To enhance robustness, bootstrapping with 1,000 iterations was applied to estimate 95% confidence intervals for key metrics such as total area (km^2^) and proportional change (%) in each suitability class. This resampling approach ensured reliable uncertainty estimation, particularly for comparisons across scenarios and suitability thresholds.

Cohen’s Kappa was calculated to evaluate the level of agreement between the observed and predicted classifications while accounting for the possibility of chance agreement.

All statistical analyses were performed in R version 4.5.0, using base functions such as `glm()` for regression modeling and additional packages including `boot` for resampling, `dplyr` for data wrangling, and `terra` for raster analysis. All procedures adhered to best practices in ecological modeling and statistical analysis to ensure methodological transparency, reproducibility, and comparability with related studies.

An analytical workflow was systematically developed using Microsoft Visio to guide the entire modeling process starting from species occurrence data acquisition and environmental variable preparation, through variable selection, ensemble species distribution modeling (SDM), projection under future climate scenarios (SSP2-4.5 and SSP5-8.5), to spatial analysis and output generation. The workflow also includes model evaluation using AUC metrics and spatially explicit post-modeling analyses such as suitability change detection and transition mapping. The complete workflow is illustrated in Fig1.tif.

**Fig 1.**
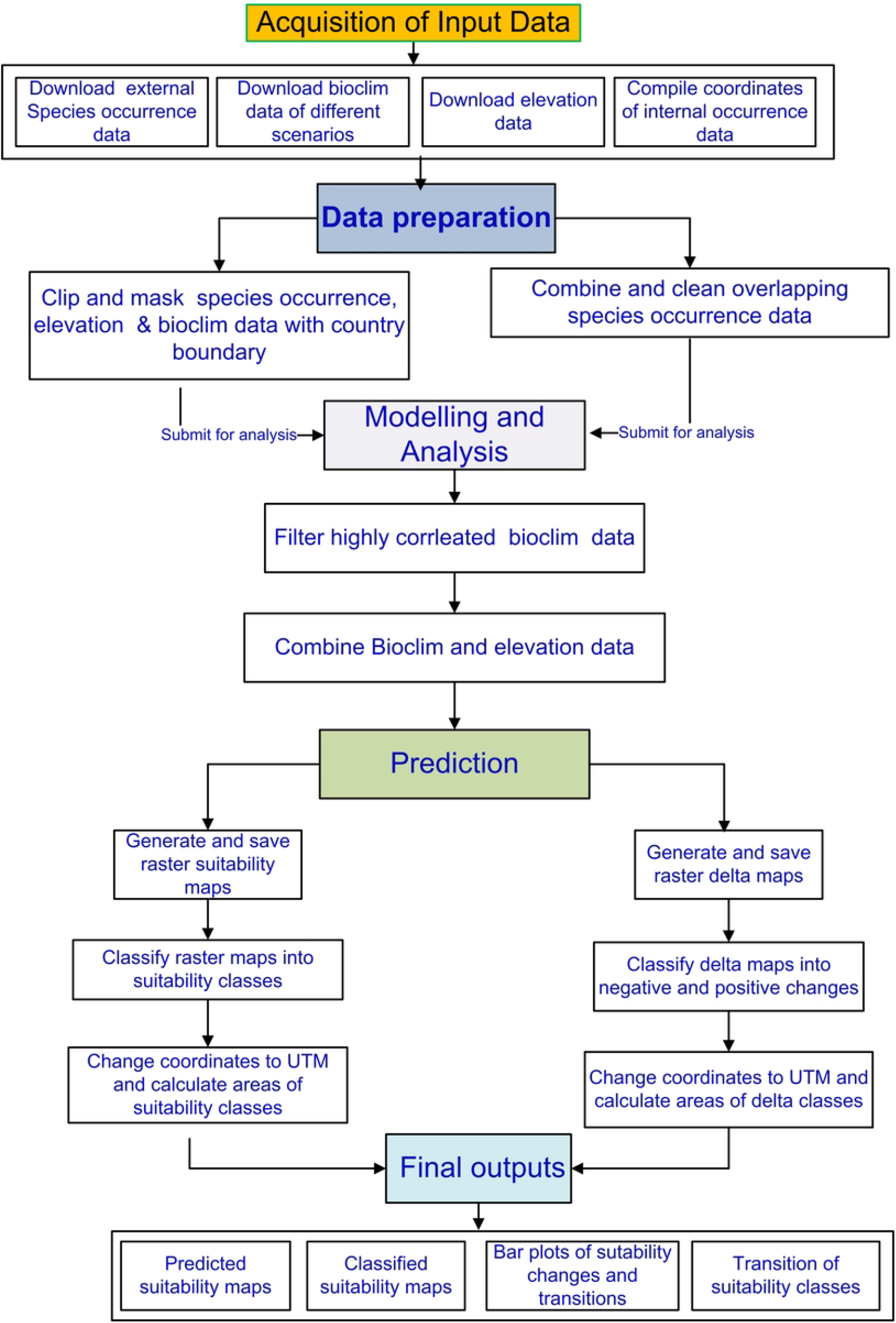
Workflow illustrating the steps of species distribution modeling (SDM) for *J*.*procera* in Ethiopia.

## 3. Results

### 3.1. Model Performance and Variable Importance

All three algorithms, Random Forest (RF), MaxEnt, and Boosted Regression Trees (BRT) treated separately and the ensemble model thereof demonstrated strong predictive performance in modeling the current distribution of *J. procera*. Our result is based on the Ensemble Model. Ensemble model performance was high, with mean AUC values exceeding 0.909 and TSS values above 0.737, indicating excellent discrimination between suitable and unsuitable habitats.

Among the environmental predictors, average temperature during the driest three-month period (BIO9), total precipitation in the driest month (BIO14), coefficient of variation of monthly precipitation (SD/mean × 100) (Bio15), total precipitation during the coldest quarter (Bio19 and elevation (to a slight degree) were consistently identified as the most influential variables across all algorithms. These variables reflect the species’ sensitivity to both moisture availability and thermal variability, particularly in the highland regions.

### 3.2. Projected Changes in Habitat Suitability under Future Climate Scenarios

The ensemble model predicted that the current suitable habitats for *J. procera* are predominantly concentrated within mid - to high-elevation zones of the Ethiopian highlands such as central and northern highlands and the southeastern highlands such as the Arsi and Bale Mountains, consistent with the known ecological niche of *J. procera*. Medium and Low suitability zones extend into adjacent elevational gradients such as arid eastern regions in the south and east as well as some parts of the Rift Valley where the species may persist under suboptimal conditions. The predicted scenario suitability maps are given in Fig2.tif. The habitat suitability map of the current scenario aligns with the known Dry Evergreen Afromontane Forest extent (Pichi Sermolli, 1957; Fiis, et al., 2010), validating the model performance.

**Fig 2.**
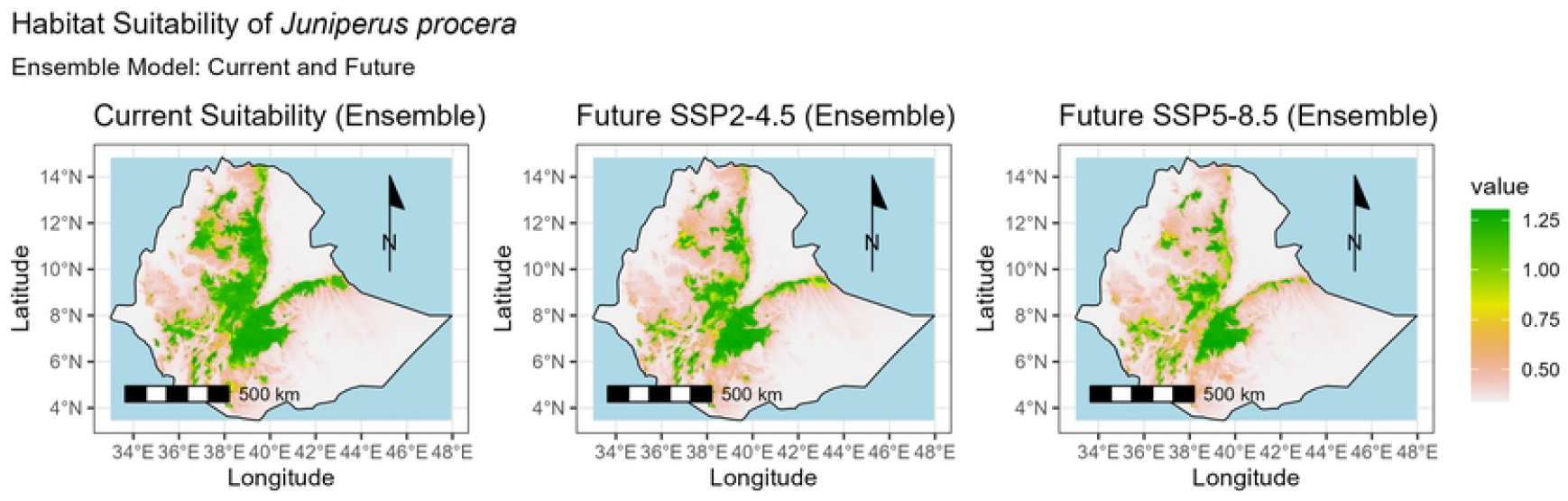
Ensemble habitat suitability predictions for *J*.*procera* under current climate and future scenarios SSP2-4.5 and SSP5-8.5. Suitability ranges from low (light pink) to high (green).

The current habitat suitability map closely aligns with the documented extent of Ethiopia’s Dry Evergreen Afromontane Forest (DAF) zones (Zerihun Woldu, 1999; Friis et al., 2010), confirming the ecological validity and spatial accuracy of the ensemble model output.

Projections from the ensemble species distribution models under both climate scenarios - moderate emissions mitigation (SSP2-4.5) and high emissions (SSP5-8.5) indicate significant spatial reconfiguration and a marked decline in highly suitable habitats for *J*.*procera* (Fig2). Under the SSP5-8.5 pathway, areas currently identified as having high habitat suitability are expected to shrink by nearly 85%, highlighting the species’ vulnerability to habitat fragmentation and range contraction. While SSP2-4.5 shows some compensatory expansion in medium and low suitability zones, particularly along the humid Afromontane highlands in the southwestern parts of Ethiopia and climate-resilient refugia in the northern region - these gains are substantially reduced under SSP5-8.5. This pattern suggests a possible downward altitudinal shift in the species’ range, likely driven by rising temperatures and altered moisture regimes, with limited highland refugia potentially serving as last strongholds for *J. procera* persistence under severe climate stress.

Under the more extreme SSP5-8.5 scenario, the remaining highly suitable habitats for *J*.*procera* are projected to be confined to a few isolated ecological refugia, primarily within the Bale Mountains in the southeastern highlands and the Gurage Mountain Range in central Ethiopia. These montane systems, which benefit from relatively cooler microclimates and persistent moisture availability, may offer temporary resilience against the escalating impacts of climate change. However, the spatial isolation and restricted extent of these habitats raise serious concerns regarding the long-term viability of *J. procera*, particularly concerning genetic connectivity, population stability, and the continued provision of its ecological functions across the broader Afromontane landscape.

To translate the continuous suitability outputs into ecologically interpretable categories, we reclassified the prediction maps into four suitability classes: High, Medium, Low, and None. This classification allowed for the spatial delineation of potential habitat zones for *J. procera*, facilitating subsequent analyses of distribution dynamics under current and future climate scenarios. The classified suitability maps are given in Fig3.tif.

**Fig 3.**
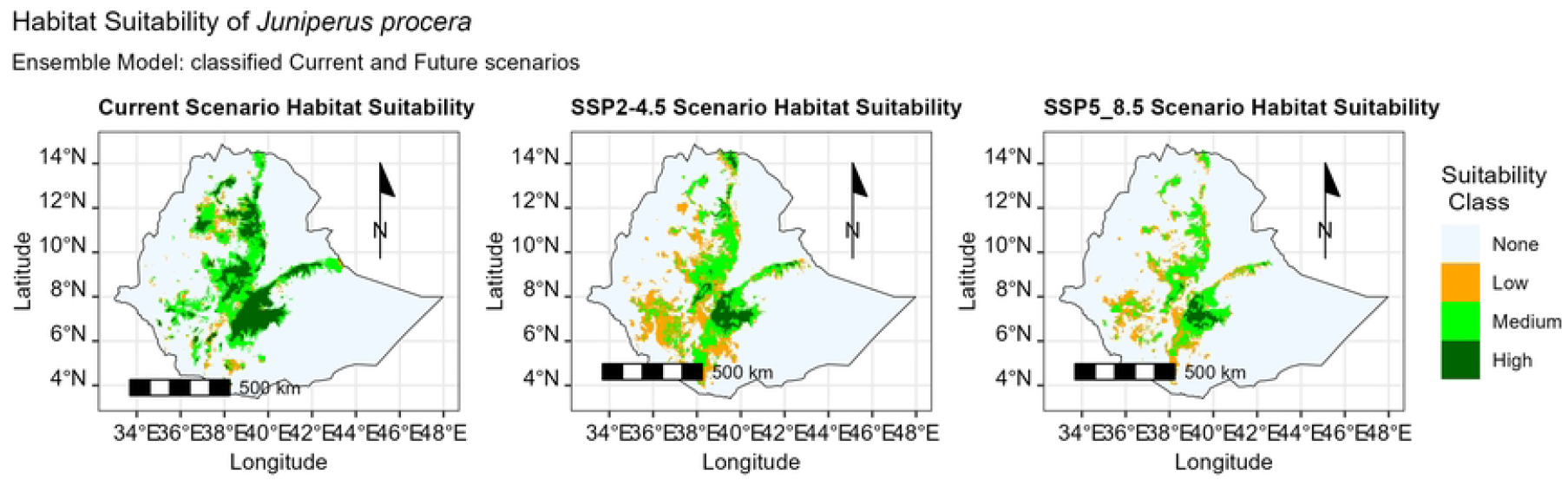
Current habitat suitability classification map for *J*.*procera* in Ethiopia, showing discrete suitability classes: None, Low, Medium, and High.

Bar plot analysis (Fig4.tif) based on the classified suitability maps quantifies these dynamics, highlighting the stark decrease in the High suitability area accompanied by expansions of Low and Medium suitability classes. These shifts underscore a general trend of habitat quality degradation under climate change, with important implications for species persistence.

**Fig 4.**
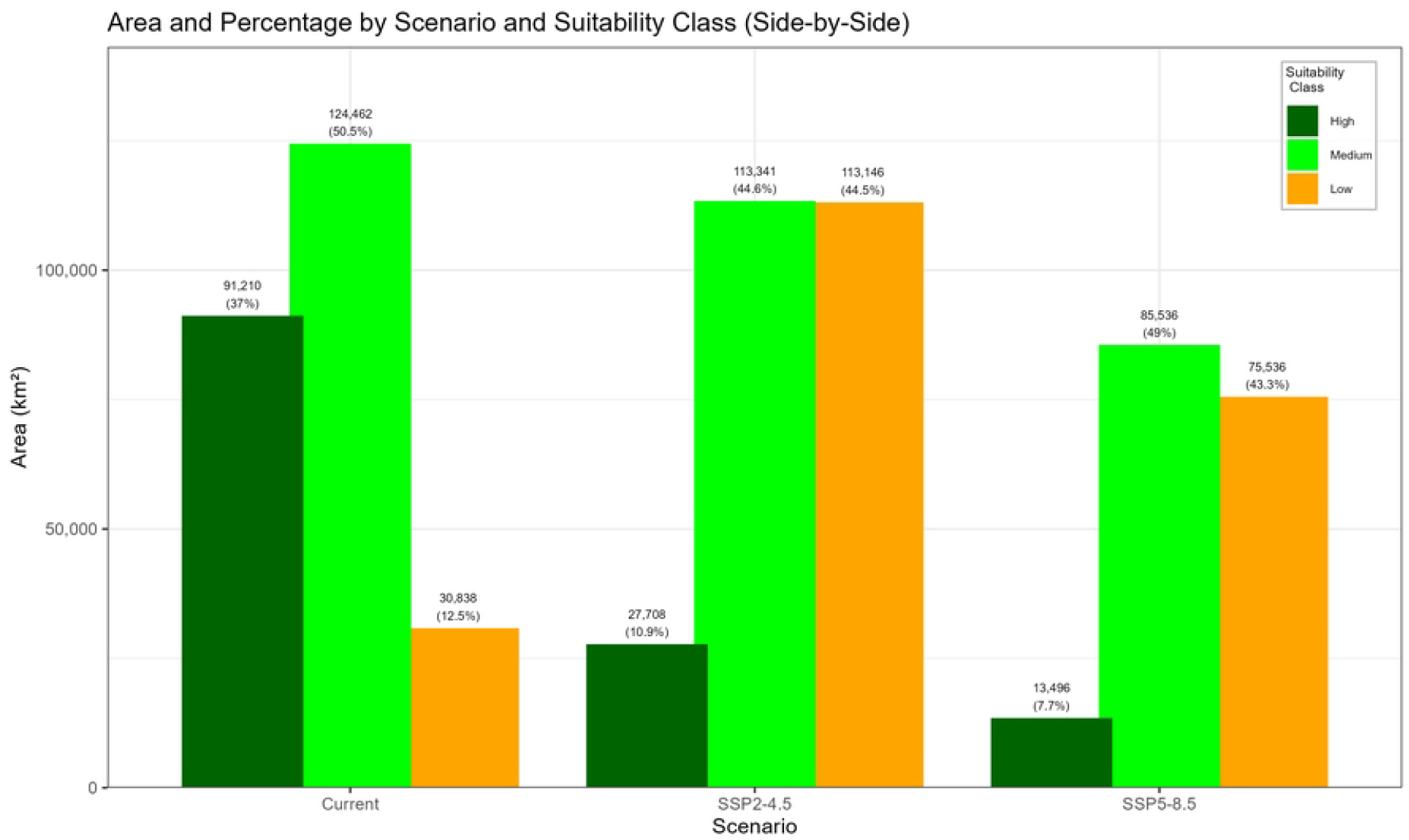
Bar plot illustrating the area (km^2^) and relative proportion (%) of habitat suitability classes (High, Medium, Low) under current and future climate scenarios.

### 3.5. Area Change Analysis and Suitability Transition

Classified delta maps (Fig5.tif and Fig6.tif) and Table 1 illustrate the spatial configuration of habitat changes for *J. procera* under both climate scenarios. Under the moderate emission pathway (SSP2-4.5), a total of 62,381 km^2^ transitioned from unsuitable (“None”) to suitable classes, with the majority shifting into the “Low” (40,693 km^2^) and “Medium” (14,480 km^2^) suitability categories. Upward transitions to higher suitability were also observed, including “Low → High” (5,750 km^2^) and “Medium → High” (5,729 km^2^), indicating potential localized improvement in habitat conditions.

**Fig 5.**
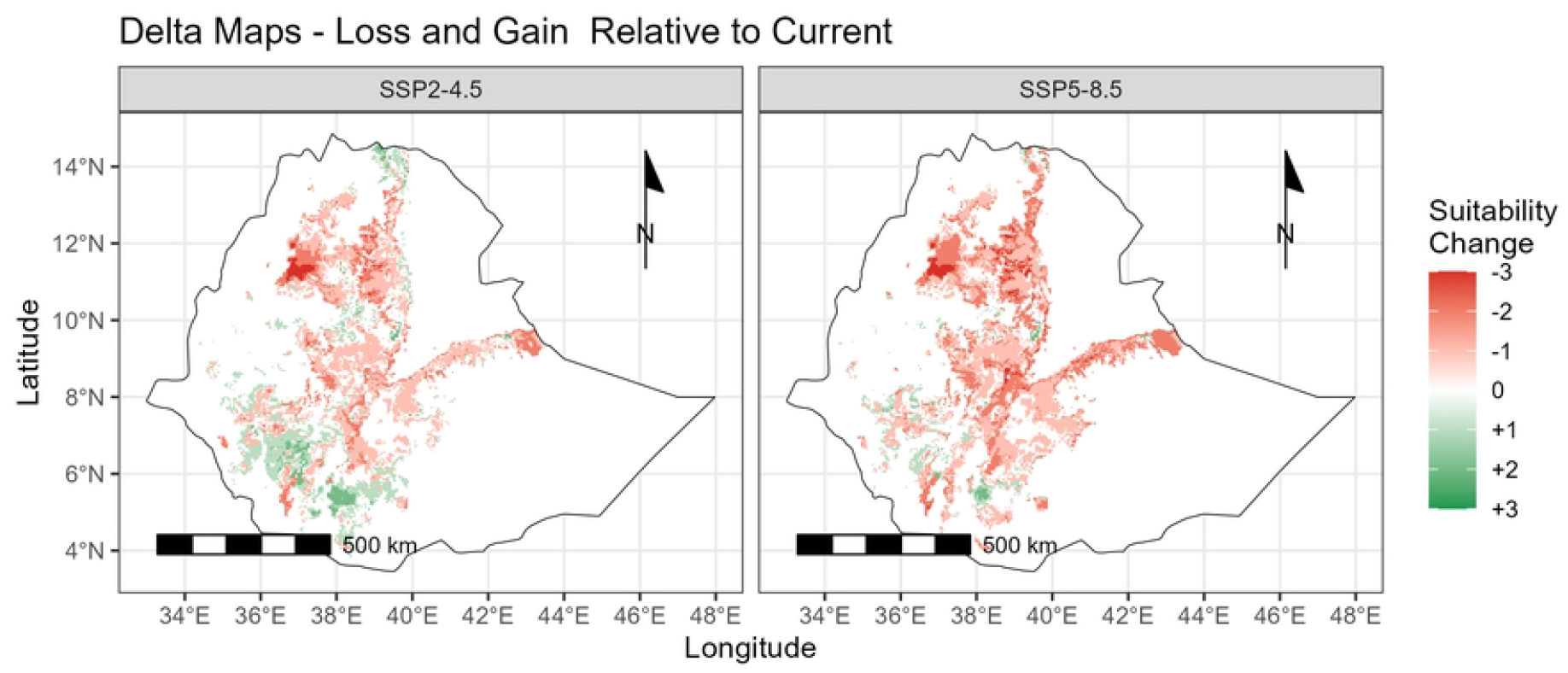
Delta maps depicting spatial changes in habitat suitability between current and future periods under SSP2-4.5 and SSP5-8.5 scenarios.

**Table 1.** Area (km^2^) of habitat suitability class gains, losses, and stable zones for *J*.*procera* under SSP2-4.5 and SSP5-8.5 scenarios.

| Scenario | Current Class | Future Class | Change Type | Area (km<sup>2</sup>) |
| --- | --- | --- | --- | --- |
| SSP2-4.5 | None | Low | Gain | 40,692.625 |
| SSP2-4.5 | None | Medium | Gain | 14,480.233 |
| SSP2-4.5 | None | High | Gain | 7,208.408 |
| SSP2-4.5 | Low | High | Gain | 5,749.815 |
| SSP2-4.5 | Medium | High | Gain | 5,728.676 |
| SSP2-4.5 | Low | Medium | Gain | 5,517.286 |
| SSP2-4.5 | High | Medium | Loss | 38,811.251 |
| SSP2-4.5 | Medium | Low | Loss | 28,939.326 |
| SSP2-4.5 | Medium | None | Loss | 16,488.440 |
| SSP2-4.5 | Low | None | Loss | 7,821.440 |
| SSP2-4.5 | High | Low | Loss | 7,652.327 |
| SSP2-4.5 | High | None | Loss | 5,496.147 |
| SSP5-8.5 | None | Low | Gain | 18,116.145 |
| SSP5-8.5 | None | Medium | Gain | 8,899.530 |
| SSP5-8.5 | None | High | Gain | 7,314.103 |
| SSP5-8.5 | Low | High | Gain | 5,348.174 |
| SSP5-8.5 | Low | Medium | Gain | 5,115.644 |
| SSP5-8.5 | Medium | High | Gain | 4,375.778 |
| SSP5-8.5 | High | Medium | Loss | 42,764.249 |
| SSP5-8.5 | Medium | None | Loss | 32,321.571 |
| SSP5-8.5 | Medium | Low | Loss | 23,295.207 |
| SSP5-8.5 | High | Low | Loss | 11,097.989 |
| SSP5-8.5 | Low | None | Loss | 11,076.850 |
| SSP5-8.5 | High | None | Loss | 6,870.183 |

**Fig 6.**
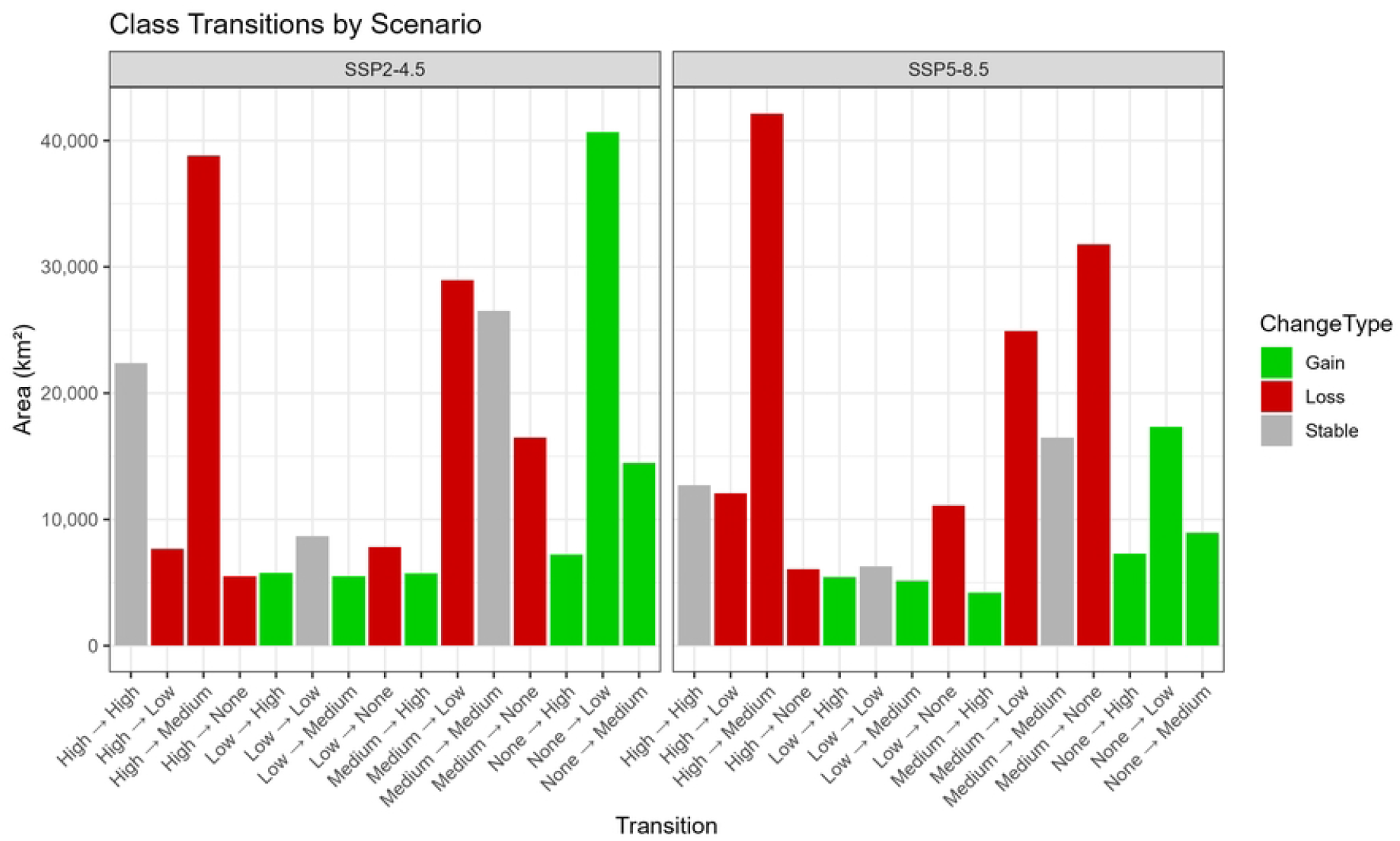
Suitability class transitions of *J*.*procera* under future climate scenarios (SSP2-4.5 and SSP5-8.5). Bars represent the total area (in km^2^) undergoing transition between classes, categorized as Gains (green), Losses (red), or Stable (not shown).

Transition matrices (Tab1; Fig6.tif.) further detail these shifts. For SSP2-4.5, although gains in suitability are evident, a notable 38,811 km^2^ of High suitability is projected to decline to Medium and 28,939 km^2^ from Medium to Low. Meanwhile, 21,984 km^2^ of previously suitable areas become unsuitable, as seen in transitions such as “Medium → None” and “High → None.” These findings suggest that even under a moderate emissions scenario, habitat degradation remains a significant concern.

Under the extreme SSP5-8.5 scenario, transitions reveal a more severe trajectory. Total gains from “None” to any suitable class decreased to 34,330 km^2^, with gains in the “Low” and “Medium” categories nearly halved compared to SSP2-4.5. While “None → High” increases slightly to 7,314 km^2^, this is offset by greater losses. Specifically, 42,764 km^2^ of High suitability is downgraded to Medium, and 32,322 km^2^ of Medium suitability transitions to None, suggesting a widespread decline in habitat quality.

Overall, the combined area transitioning from any suitable class to unsuitability under SSP5-8.5 exceeds 50,000 km^2^, underscoring the severity of habitat contraction for *J. procera* under high-emissions conditions. These trends highlight the growing vulnerability of the species’ climatic niche and the limited capacity for natural range compensation in the face of intensifying climate pressures.

### 3.6. Statistical Significance of Habitat Changes

The binary classification results of the species distribution model demonstrated a substantial level of agreement with observed presence-absence data, as reflected by a Cohen’s Kappa value of 0.734. This indicates that the model performed well in distinguishing suitable from unsuitable habitats, even after correcting for chance agreement. Furthermore, generalized linear modeling (GLM) and Chi-squared tests provided strong statistical support for the influence of climate scenarios on habitat distribution. The GLM results showed a highly significant effect of climate scenario on the distribution of habitat suitability classes (p < 0.001), while the Chi-squared test revealed an even more pronounced significance (p < 2.2 × 10^−16^). Together, these findings reinforce the conclusion that future climate trajectories will drive major shifts in the spatial configuration of suitable habitats for *J. procera*.

## 4. Discussion

Our ensemble species distribution models (SDMs) forecast substantial shifts in the spatial configuration and ecological quality of *J. procera* habitat across Ethiopia under both moderate (SSP2-4.5) and high-emission (SSP5-8.5) scenarios. The conversion of continuous suitability scores into discrete classes revealed critical patterns of habitat loss, gain, and internal reorganization, enabling a nuanced interpretation of climate-driven range dynamics.

A dominant pattern emerging from our projections is the contraction and fragmentation of high-suitability zones, particularly under SSP5-8.5, where climatically suitable habitat of *J. procera* from 328,819 km^2^ under current conditions to 174,568 km^2^ under SSP5-8.5 - a reduction of 47% and more than 42,700 km^2^ (approximately 85%) of currently high-suitability habitat transitions to lower suitability categories or becomes climatically unsuitable. Even under SSP2-4.5, moderate emissions are projected to drive significant declines, with over 55,000 km^2^ of medium and high-suitability habitat degraded or lost. These findings corroborate global evidence that montane and Afromontane species are especially vulnerable to climate change due to their narrow climatic niches and limited upslope migration capacity (Luedeling et al., 2011; Morán-Ordóñez et al., 2017).

This projected habitat degradation poses serious implications for *J. procera*, a slow-growing, long-lived conifer with low recruitment rates and narrow physiological tolerance (Adams, 1990; Teketay & Granström, 1995; Gebirehiwot et al., 2024). As a canopy-dominant and ecologically pivotal species, *J. procera* supports a wide array of flora and fauna, contributes to microclimatic regulation, and stabilizes soil and hydrological systems (Paine, 1969; Zerihun Woldu, 1999). Declines in its distribution are therefore likely to initiate cascading ecological effects, including biodiversity loss and ecosystem service disruption.

Further compounding the threat is the pronounced spatial fragmentation of remaining suitable habitats, particularly in highland regions where upslope migration is constrained by topography and land-use change (Negash, 2010). Fragmentation may isolate populations, inhibit gene flow, and reduce the species’ capacity to adapt to future environmental variability—a concern also raised in studies of other Afromontane taxa (Borghesio et al., 2004).

Despite these declines, our models also predict the emergence of climatically suitable habitats in previously unsuitable areas, especially along wetter areas in the southwest and higher elevation zones. Under SSP2-4.5, over 62,000 km^2^ of new suitable habitat is projected to emerge, including approximately 7,200 km^2^ of high suitability. These newly suitable areas may serve as potential refugia if colonization and establishment are feasible. However, model-based suitability does not guarantee species persistence; real-world establishment depends on propagule pressure, dispersal limitations, soil conditions, and interspecific interactions, none of which are captured by correlative SDMs (Guisan et al., 2017; Franklin, 2010). In Ethiopia, historical land degradation, soil nutrient depletion, and the collapse of seed banks further constrain natural regeneration (Gete Zeleke & Hurni, 2001).

Observed transitions within currently suitable zones - from low to medium or medium to high - offer some positive signals, suggesting potential for local improvement in habitat quality under specific scenarios. These internal gains may reflect microrefugia, which have been shown to support persistence in Mediterranean and montane ecosystems despite broader climatic pressures (Araújo & New, 2007).

Importantly, habitat shifts projected for *J. procera* mirror wider ecological trends in the Ethiopian highlands. Several studies have documented climate-induced altitudinal shifts, particularly the upward expansion of drought-adapted C_4_ grasses into formerly C_3_-dominated ecosystems (Zerihn Woldu, 1985, 1991; Still et al., 2003). These shifts reflect biome-level reorganizations under rising temperature and declining moisture regimes (Sage & Monson, 1999), and further underscore the vulnerability of endemic Afromontane flora to climate stress.

In Ethiopia, these climatic risks are compounded by severe anthropogenic pressures including deforestation, fuelwood harvesting, overgrazing, and agricultural encroachment (Bewket & Sterk, 2005; Teketay, 1992; Aerts et al., 2016). These land-use changes have fragmented natural forests, disrupted regeneration dynamics, and intensified erosion—thereby amplifying the impact of climate change on already stressed ecosystems (Hurni et al., 2015; Tolessa et al., 2017).

As a keystone species, *J. procera* serves as a biological indicator of ecosystem integrity. The 85% loss of high-suitability area alone represents a profound decline in the species’ optimal niche space, with likely consequences for its long-term viability. Its contraction under future scenarios may therefore signal broader degradation of Afromontane forest systems and loss of associated ecological functions. These findings underscore the urgency of conservation interventions that integrate climate adaptation and ecological restoration.

From a management perspective, our results highlight critical spatial targets for conservation and restoration. Projected refugia and persistent high-suitability zones should be prioritized for protection, particularly those overlapping with Ethiopia’s national restoration programs such as the Green Legacy Initiative. These areas offer strategic opportunities to reinforce ecological networks, enhance landscape connectivity, and facilitate upslope migration where possible. Assisted migration and enrichment planting may be required to establish populations in newly suitable zones with limited natural dispersal.

## 5. Conclusion

This study highlights the acute vulnerability of *J*.*procera* to future climate change across its native Ethiopian range. Using ensemble species distribution models (SDMs), we project a substantial contraction of high-suitability habitats under both moderate (SSP2-4.5) and high-emissions (SSP5-8.5) scenarios. Under SSP5-8.5, habitat classified as highly suitable is expected to decline by approximately 85%, posing a significant threat to the long-term persistence of this keystone Afromontane species.

Transition analysis reveals a complex spatial redistribution, including potential gains in suitability within areas currently deemed unsuitable. These newly emerging zones often in lower elevation or mid-altitude regions suggest a possible downward altitudinal shift in contrast to the more commonly anticipated upslope migration. However, such gains must be interpreted with caution. Many of these areas may lack the soil conditions, hydrology, and biotic interactions necessary to support successful establishment and regeneration. The abrupt transitions from “None” to “High” suitability, as modeled here, may reflect rapid climatic shifts rather than ecologically viable colonization potential, increasing the risk of bottlenecks, fragmentation, and localized extinctions.

These dynamics point to a broader ecological reshuffling, with simultaneous loss of core habitats and the emergence of climatically suitable but ecologically uncertain zones. Without timely and integrated conservation strategies, *J. procera* is likely to experience range contraction, fragmentation, and increasing marginalization within its native ecosystem. Moreover, if global climate change continues unabated and land degradation remains unaddressed, the combined ecological and social consequences could escalate beyond manageable thresholds. Proactive, targeted interventions are urgently required to reduce vulnerability, enhance species adaptability, and maintain the ecological functions and services that support both biodiversity and human livelihoods.

## 6. Recommendations

Based on the findings of this study, we recommend the following actions for conservation planning, policy formulation, and climate adaptation strategies:

1. Protect Climatic Refugia Immediate protection should be prioritized for areas currently suitable and projected to remain stable under future climate scenarios. These refugia act as critical strongholds for *J. procera*, preserving genetic diversity and population resilience.
2. Restore Emerging Yet Degraded Habitats Areas projected to become suitable but currently degraded should be rehabilitated through reforestation, soil conservation, and erosion control. Proactive restoration in these transition zones will improve colonization potential and ecosystem readiness.
3. Enhance Landscape Connectivity Establish ecological corridors and stepping-stone habitats to facilitate migration and gene flow. This is especially important in topographically complex regions where natural dispersal is limited by elevation and fragmentation.
4. Implement Assisted Migration Where Necessary In highly fragmented landscapes or distant newly suitable areas, assisted migration should be considered. This may involve nursery propagation, seedling translocation, or community-led planting initiatives in climatically favorable but unoccupied habitats.
5. Integrate SDM Projections into Forest Policy National and regional afforestation programs such as Ethiopia’s Green Legacy Initiative should incorporate SDM-based forecasts to ensure alignment with future climate suitability rather than historical distributions alone.
6. Establish Long-Term Monitoring Programs Permanent ecological monitoring plots across elevational gradients should be developed to track population dynamics, recruitment rates, and community shifts over time, informing adaptive management.
7. Engage Local Communities and Institutions Conservation success hinges on the active participation of local stakeholders. Incentive-based conservation, integration of traditional ecological knowledge, and environmental education programs should be embedded in planning frameworks.
8. Promote Policy Integration and Transdisciplinary Research Effective response to climate-driven habitat shifts requires collaboration across forestry, climate science, ecology, and socioeconomics. Future research should combine biophysical modeling with institutional and social analyses to identify practical, community-supported conservation strategies.

## 7. Data Availability

All occurrence data, environmental predictor layers, and R scripts utilized in this study can be obtained from the corresponding author upon reasonable request. The ensemble modeling scripts, along with additional tools for classifying predicted raster outputs, calculating area, generating delta maps, and analyzing transition dynamics, were compiled and further refined by the corresponding author.

GBIF data are publicly accessible via the GBIF portal (https://www.gbif.org), with specific query details documented in the supplementary materials.

Climate data (current and future bioclimatic variables) were obtained from WorldClim version 2.1 (https://worldclim.org) and CMIP6 repositories, with details on model selection, scenario choice, and downscaling procedures fully documented for transparency. Elevation data were also obtained from the same source.

Processed spatial data products and model outputs generated in this study will be archived in an open-access repository upon publication to facilitate reuse and further research.

## 9. Statements and Declarations

All authors contributed to the study conception and design. Material preparation, data collection and analysis were performed by Zerihun Woldu. Gete zeleke also participated in material preparation. The first draft of the manuscript was written by Zerihun Woldu. Gete Zeleke commented on previous versions of the manuscript and approved the final manuscript.

We have not received funds, grants, or other support during the preparation of this manuscript. We have no financial or non-financial interests that are directly or indirectly related to this work.

## Declaration of originality

The authors confirm that this manuscript has not been published, is not under consideration for publication, and has not been submitted elsewhere, either in part or in whole. It is an original work prepared exclusively for submission to *Plos One*.

## Notes

### Competing Interest Statement

The authors have declared no competing interest.

